# Mechanisms of chromatin remodeling by an Snf2-type ATPase

**DOI:** 10.1101/2024.12.31.630910

**Authors:** Deepshikha Malik, Ashish Deshmukh, Silvija Bilokapic, Mario Halic

**Affiliations:** Department of Structural Biology, St. Jude Children’s Research Hospital, 262 Danny Thomas Place, Memphis, TN, 38105, USA

**Keywords:** SNF2H, chromatin remodeling, ISWI, nucleosome, cryo-EM, remodelers

## Abstract

Chromatin remodeling enzymes play a crucial role in the organization of chromatin, enabling both stability and plasticity of genome regulation. These enzymes use a Snf2-type ATPase motor to move nucleosomes, but how they translocate DNA around the histone octamer is unclear. Here we use cryo-EM to visualize the continuous motion of nucleosomal DNA induced by human chromatin remodeler SNF2H, an ISWI family member. Our work reveals conformational changes in SNF2H, DNA and histones during nucleosome sliding and provides the structural basis for DNA translocation. ATP hydrolysis induces conformational changes in SNF2H that pull the DNA tracking strand, distorting DNA and histones at SHL2. This is followed by SNF2H rotation on the nucleosome, which first pulls the DNA guide strand and creates one-base pair bulge at SHL2, and then releases the pulled DNA. Given the high conservation of the catalytic motors among ATP-dependent chromatin remodelers, the mechanisms we describe likely apply to other families.

---

In eukaryotes, ATP-dependent chromatin remodeling enzymes enable genome regulation and maintenance by actively shaping the packaging of genomic DNA. Chromatin remodelers regulate DNA accessibility and thereby genome transactions such as transcription, DNA replication and DNA repair. Nucleosomal DNA translocation by remodelers alters and disrupts histone–DNA contacts, allowing DNA mobilization around histones (‘nucleosome sliding’), histone ejection and exchange with histone variants or unmodified histones. During nucleosome sliding by remodelers, DNA is translocated from the entry site, around the histone octamer, and emerges at the exit site ^1–3^.

Chromatin remodelers are helicase-like motor enzymes, classified into four families: SWI/SNF, ISWI, INO80 and CHD. All remodeler enzymes share a Snf2-type ATPase domain, composed of two conserved RecA-like lobes that form the ATPase site. The two lobes bind both DNA strands and are sufficient to couple ATP hydrolysis to chromatin remodeling. A C-terminal segment forms the so-called brace helix, which spans across both lobes and couples their motions ^1,4–6^.

The interactions between chromatin remodelers and nucleosomes have been extensively studied, but most cryo-EM structures have been determined using samples that were chemically cross-linked before freezing, which limits any dynamics that occur during chromatin remodeling ^7–13^. Several of those structures showed remodelers in different nucleotide-bound states ^11–15^. In particular, Snf2 and ISWI were captured bound to non-hydrolyzable ADP-BeFx, to ADP, or in their nucleotide-free forms, providing snapshots of the DNA translocation cycle. The ADP-BeFx-bound structures showed the nucleosome in a near-canonical conformation, indicating that they represent a state prior to ATP hydrolysis and DNA translocation ^11,12,15^. In contrast, the ADP-bound structures showed a 1-bp bulge in the nucleosomal DNA at superhelix 2 (SHL2), suggesting they represent a post-hydrolysis state ^11,12,16^. Moreover, in the ADP-bound structures, the DNA tracking strand (i.e. the one topologically equivalent to the single strand on which DNA or RNA helicases translocate) showed a distortion from the entry site to SHL2 ^11,12^. This latter observation suggests that remodelers operate through an asynchronous mechanism for DNA translocation, in which the DNA strands translocate at different times, but no putative intermediate states have so far been captured ^4–6^.

Studies using NMR and cysteine cross-linking approaches have shown that binding of human SNF2H to the nucleosome deformed the histone octamer, and those distortions were essential for remodeling activity ^17^. However, such histone alterations have not been observed in the currently available cryo-EMstructures of nucleosome-remodelers ^11,12,15,18,19^. It is plausible that those structures represent ground states, stabilized by the presence of ADP or ADP-BeFx, or by cross-linking, and they thus could not capture alterations to the nucleosome driven by ATP hydrolysis. This limitation leaves many open questions regarding the mechanism for nucleosome sliding by remodelers. How exactly is DNA translocated during nucleosome sliding? How is directionality achieved? Does nucleosomal DNA movement induce changes in histones?

To address these gaps, we present a set of cryo-EM structures of human SNF2H actively remodeling nucleosomes. Our data reveal how SNF2H moves DNA on the nucleosome and show the conformational changes in SNF2H, DNA and histones during DNA translocation. As the catalytic motors of chromatin remodelers are highly conserved, the mechanisms we describe for SNF2H likely apply to other families of ATP-dependent chromatin remodeling enzymes.

## RESULTS

### SNF2H binding to nucleosome

We purified recombinant human SNF2H and reconstituted a nucleosome with a DNA fragment containing the 601 sequence ^20^ and 80bp of linker DNA on one side (**Figure S1a**). The purified SNF2H was able to bind to the nucleosome (**Figure S1b**) and to remodel it upon activation by ATP (**Figure S1c**). To investigate the mechanisms of DNA translocation, we collected cryo-EM images of the ATP-actived complex frozen at multiple time points (5s, 2min and 10min) after addition of ATP and with two different concentrations of MgCl_2_ (**Figure S1d**). We merged the data from all conditions and performed extensive data analyses to determine 13 unique structures of the active SNF2H bound to the nucleosome (**Figure S1e-i, 2a and Table 1**). The datasets collected under different Mg2+ concentrations or time points after activation showed the SNF2H-nucleosome complex in the same conformations, with small variations in the fraction of particles belonging to each conformation (**Figure S1j**), likely because the complex undergoes multiple cycles of DNA translocation that are not synchronized prior to being frozen. Moreover, the enzyme is found on different DNA sequences found at SHL2 at any given time point, indicating that the mechanisms of DNA translocation do not seem to be sequence-specific. In agreement with previous structures ^11,15,21^, SNF2H binds to the nucleosome at SHL2 (**Figure S2a**). In each structure, the nucleosome has clearly resolved amino acid side chains and DNA bases, which allowed us to build precise models for the histones and DNA (**Figure S2b**); the flexible linker DNA region was not observed. The density corresponding to SNF2H ATPase motor domain has visible side chains in most conformations, allowing us to precisely model them (**Figure S2b**). We also observed density for a nucleotide in the active site of SNF2H in nearly all conformations (**Figure S3a**).

**Table 1:**
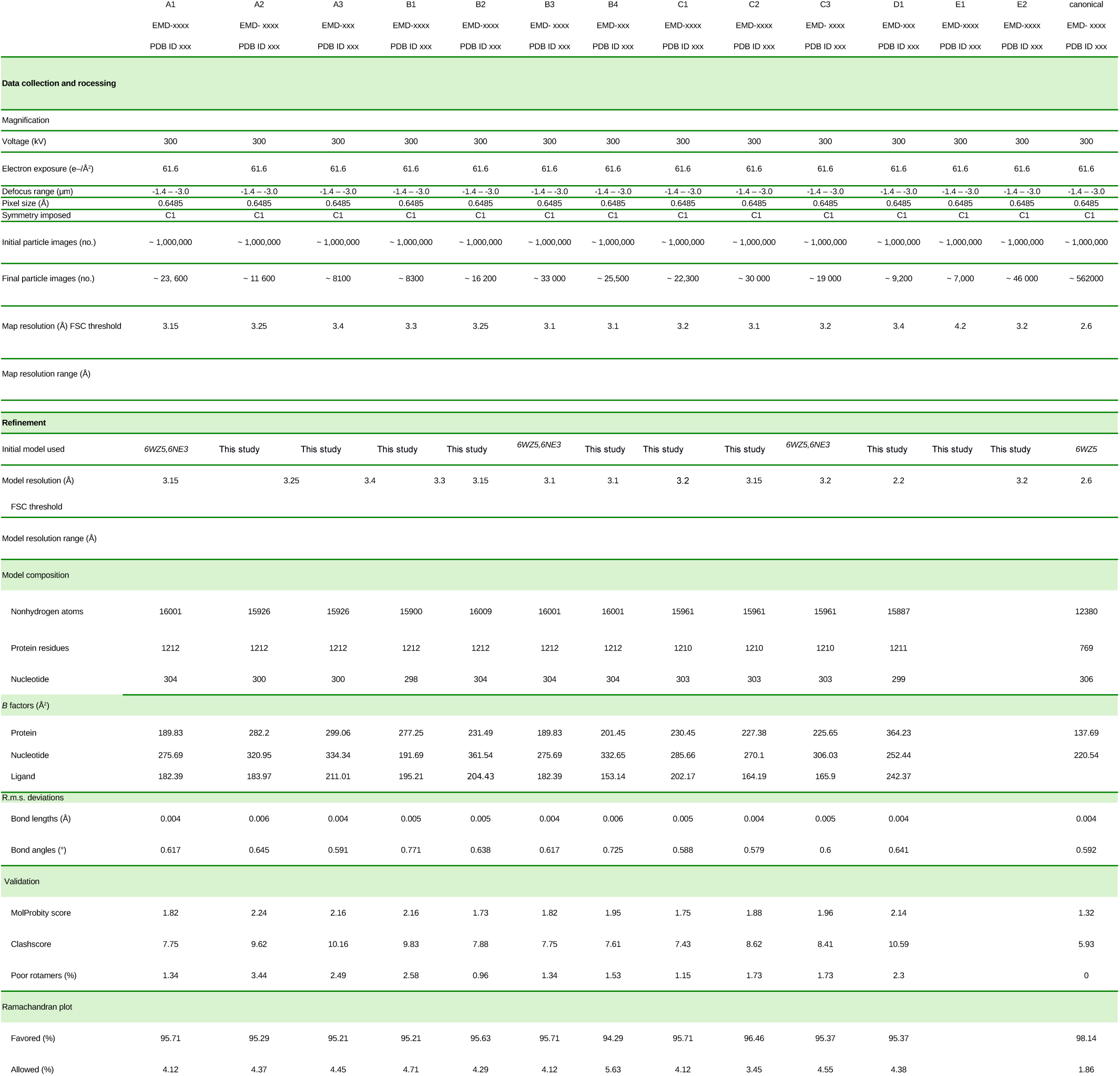
Cryo-EM data collection, refinement, and validation statistics.

Based on the DNA conformations at SHL2 and SNF2H conformations, we clustered our 13 SNF2H-nucleosome structures into five groups (A to E) (**Figure 1 and S3b-c**). In groups A to D, all components are present at high resolution: in group E, SNF2H is flexible and at low resolution. Since DNA translocation is a directional movement, we were able to establish their order relative to each other (**Figure S3b-c**). In group A (three structures, ATP hydrolysis), the DNA adopts a conformation overall similar to that in the structures of nucleosome-remodeler complexes with ADP-BeFx, i.e with the DNA slightly pulled out at SHL2, away from the histone octamer surface. In group B (four structures, tracking strand movement), the DNA tracking strand is being pulled out more than the guide strand, which deforms the double helix locally at SHL2 and SHL3. In group C (three structures, guide strand movement), the guide strand is also pulled out, correcting the DNA distortion seen in group B and allowing the double helix to accommodate one additional base pair at SHL2. In group D (one structure, guide strand release), the DNA starts moving back toward the nucleosome surface. In group E (two structures), the DNA is not bound to SNF2H, which is flexibly bound to nucleosome.

**Figure 1.**
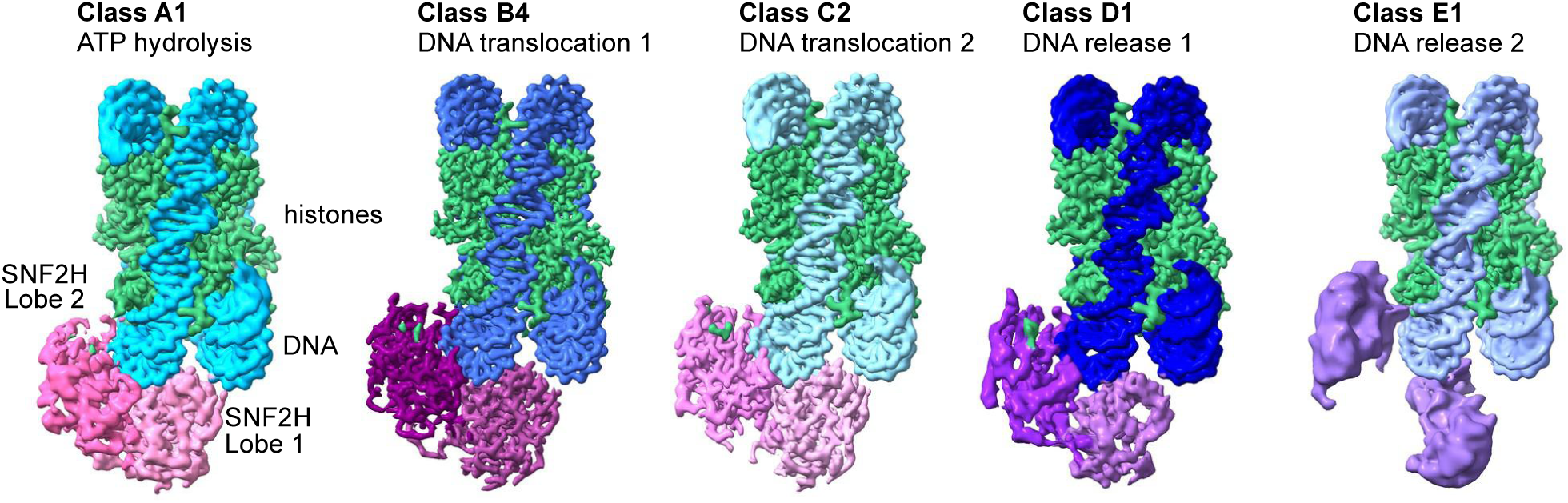
Cryo-EM structures of active human SNF2H bound to the nucleosome. Cryo-EM maps show the nucleosome-SNF2H complex during ATP hydrolysis (class A1), tracking strand movement (class B4), guide strand movement (class C2), DNA release state 1 (class D1), and DNA release state 2 (class E1). SNF2H is shown in shades of pink/purple; DNA is shown in shades of blue; histones are shown in green.

These changes in DNA conformation are coupled to structural alterations in SNF2H, and the changes in both DNA and SNF2H define the different classes within each group, as we describe below.

### ATP hydrolysis

In the group A structures (A1 to A3, 15% of A-D particles) (**Figure S1j**), we observe gradual rotation of SNF2H lobe 2 towards SHL3, relative to the structures of nucleosome-SNF2H with ADP-BeFx ^15^ (**Figure S4a-b**). The first structure (A1) shows SNF2H in a conformation overall similar to that in the previous structures of SNF2H ^15^ or ISWI ^11^ bound to ADP-BeFx, except for the rotation of SNF2H lobe 2 (**Figure S4a-b**). The DNA at SHL2 is pulled out by 1–2 Å relative to the canonical nucleosome structure (**Figure S4c-d**), which is similar to the position of the DNA in the structure of nucleosome-SNF2H with ADP-BeFx ^15^. The density of the nucleotide bound to SNF2H is in a position similar to that of ADP-BeFx in the previous structures ^15^ (**Figure S4e**) and is consistent with occupancy of either ATP or ADP-Pi. In the A2 and A3 structures, we observe further rotation of the lobe 2 towards SHL3, compared to the A1 structure (**Figure S4f-g**). These differences indicate that the group A structures represent conformational states downstream of the ADP-BeFx-bound state.

### Tracking strand movement

In the four group B structures (B1 to B4, 45% of A-D particles) (**Figure S1j**), the density for SNF2H and bound nucleotide resemble that in the previous structures of remodelers with bound ADP ^11,12^, indicating that these structures correspond to post-ATP hydrolysis states (**Figure S5a**). SNF2H adopts a canonical open conformation, identical to that of ADP-bound ISWI (**Figure S5b**), but its position relative to the nucleosome is different from the ADP-bound structures (**Figure S5a**). Compared to group A structures, SNF2H lobe 2 moves further toward SHL3, while the DNA at SHL2 is pulled out more and moves toward the dyad, in the opposite direction of lobe 2’s movement. For example, compared to A3, in the B1 structure the SNF2H lobe 2 continues to rotate toward SHL3, whereas the brace helix and lobe 1 remain stable (**Figure S5c-d**). The other group B structures show gradual conformational transitions in SNF2H: relative to the B1 structure, in the B2–B4 structures, SNF2H moves further outward, which leads to movement of the brace helix away from the nucleosome (**Figure S5e-f**).

By overlaying the A1 and B4 structures, we can visualize the extent of lobe 2 rotation toward SHL3 and the outward rotation of the brace helix (**Figure 2a**). Lobe 1 rotates in the same direction as lobe 2, but its rotation is more limited (**Figure S5g**), and the interactions between guide strand and lobe 1 or brace helix are not altered (**Figure 2b**). In contrast, the movements of lobe 2 alter some of the interactions with the DNA tracking strand (**Figure 2b**). For example, T509 interacts with the nucleotide at position −21 in group A and at position −22 in group B structures (nucleotide numbering is relative to the dyad; negative numbers mark the tracking strand). Likewise, R538 interacts with the nucleotide at −18 in A and at −19 in group B structures. Some lobe 2 residues interact with the same nucleotide in both group A and B structures, such as K455 with nucleotide at −22. We assessed the importance of the lobe 2 interactions with the tracking strand by mutating R538 or K455 to Ala. The single mutations severely reduced the chromatin remodeling activity of SNF2H (**Figure 2c**), while showing a small impact on its ATPase (**Figure S5h**) or nucleosome binding activity (**Fig S5i**). Similar results were obtained with the double mutant (K455A R538A) (**Figure S5h-j**). These observations support the importance of those interactions for DNA translocation.

**Figure 2.**
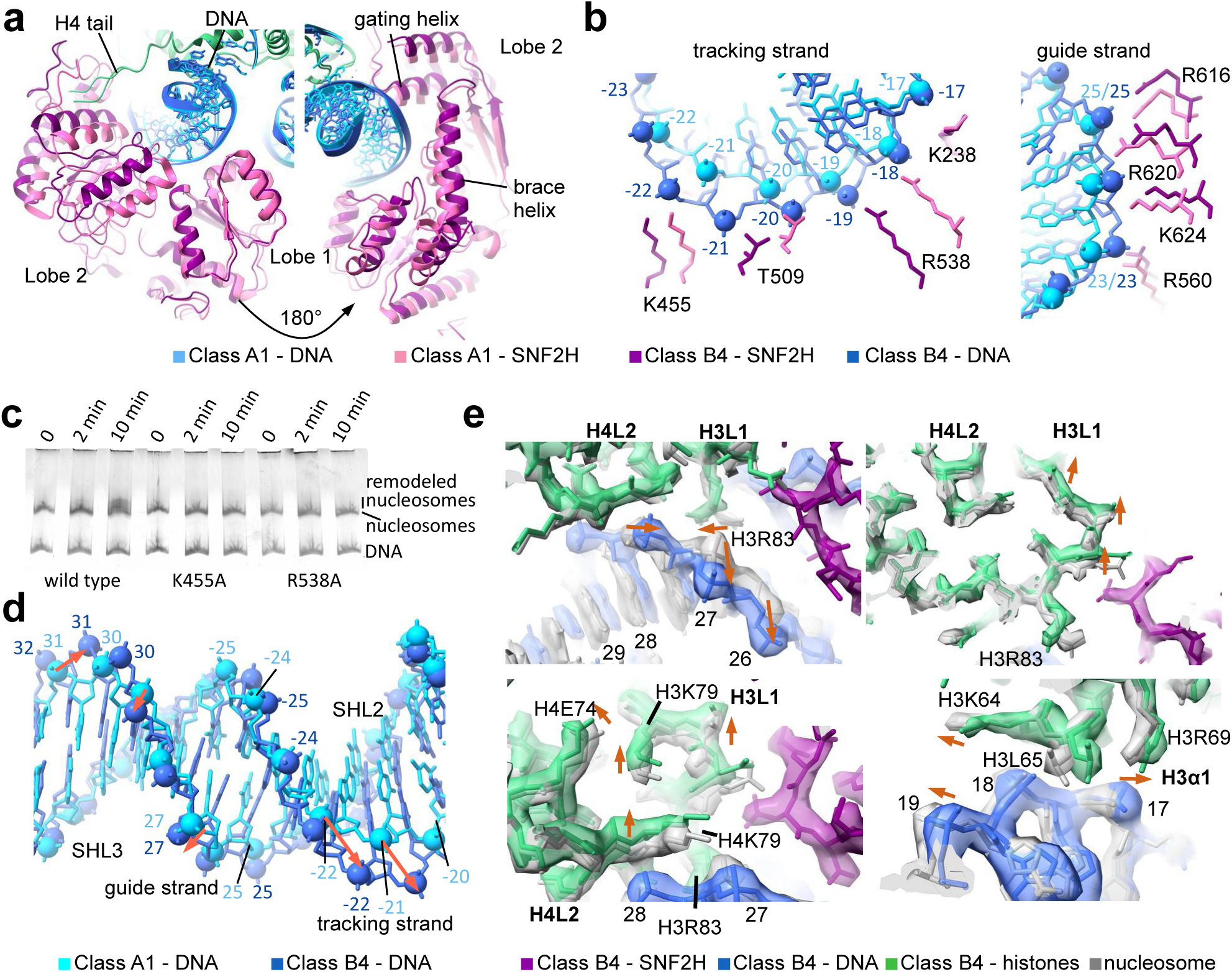
Guide strand translocation. **a)** Close-up views of the overlay of class B4 (purple) and A1 (pink) structures, showing conformational changes in SNF2H. **b)** Close-up views of the overlay of class B4 (purple, royal blue) and A1 (pink, light blue) structures, showing changes in interactions between SNF2H and DNA. Nucleotide numbering is relative to the dyad; negative numbers mark the tracking strand, positive numbers mark the guide strand. **c)** Native gel stained for DNA showing nucleosome remodeling by wild-type and K455A or R538A mutant SNF2H. **d)** Overlay of the nucleosomal DNA at SHL2 and SHL3 in class B4 (dark blue) and A1 (light blue) structures. The tracking strand is pulled out more than the guide strand, leading to DNA distortion and tilting of the bases. DNA phosphate group movements are indicated by orange arrows. **e)** Close up views of the overlay of class B4 (colored) and the canonical nucleosome (grey) structures, showing changes in the histone residues interacting with DNA at SHL2 and SHL3. Histone residue movements are indicated by orange arrows.

The movement of SNF2H lobe 2 in the group B structures pulls the nucleosomal DNA away from the histone octamer surface at SHL2 and SHL3. Both guide and tracking DNA strands are displaced outward in B1 compared to the A3 structure, but to different extents: the tracking strand is pulled out by 5-6 Å, while the guide DNA strand is pulled out by ∼1–2 Å in B1 structure (**Figure S6a**). This movement of the guide DNA strand occurs gradually, and the guide strand is pulled by ∼3–4 Å in the B4 structure (**Figure 2b, d and S6b-d**). From the entry site up to SHL3, both strands of DNA adopt the canonical conformation but they advance by 1 bp compared to their initial position in the canonical nucleosome (**Figure S6e**). At SHL2 and SHL3, only the tracking strand is pulled by one nucleotide (**Figure 2d**), as sliding of the guide DNA strand is blocked by the brace helix, which only partially moves outward in B4 structure (**Figure 2a, b**). The dissimilarity in the movements of the tracking and guide strands results in deformation of the DNA backbone and tilting of the bases at SHL2 and SHL3, which is necessary to accommodate the 1 nt difference between guide and tracking strand (**Figure 2d and S6c-d)**.

### Histone adaptation to DNA movement

As the brace helix moves outward from B1 to B4, the guide strand DNA is gradually pulled, and in B4 the DNA phosphate groups and bases are in an intermediate position, in-between two canonical positions (**Figure S6f**). To accommodate the partial movement and local distortion of the nucleosomal DNA, histones H3 and H4 undergo rearrangements in the group B structures compared to the canonical nucleosome. These rearrangements are most pronounced in the B4 structure (**Figure 2e and S6g**). In particular, residues in loops H3 L1 (E76–S86) and H4 L2 (A76–T80), which interact with DNA at SHL2.5, show several structural alterations. For example, in the B4 structure the nucleotide 28 is in a position between those of nucleotides 27 and 28 in the canonical nucleosome structure. This DNA change induces an inward flipping of H3R83, which interacts with DNA phosphate at position 28 in B4 structure, instead of position 27 as observed in group A or canonical nucleosome structures (**Figure 2e**). The DNA phosphate at position 28 moves closer to the histones and pushes H4K79 toward the histone core. Thus, accommodations to DNA changes lead to movement of the entire H3 L1 and H4 L2 loops, also affecting residues that do not interact with DNA, such as H4E74 or H3K79. We also observe rearrangements of H3K64 and H3L65 side chains in H3 α1 helix, which interacts with DNA at SHL1.5; and of H4E27 and H4E52 in H4 α1 and α2, adjusting to the movement of the H4 tail (**Figure S6g**). Importantly, on the nucleosome side where SNF2H is not bound, we do not observe any conformational changes in the histone octamer (**Figure S6h**), supporting our conclusion that these movements are due to SNF2H’s remodeling activity. To test the importance of histone adaptation for nucleosome remodeling, we non-specifically cross-linked histones with glutaraldehyde (**Figure S6i, j**) and observed that SNF2H is unable to mobilize nucleosomes containing cross-linked histones (**Figure S6k**).

### Guide strand movement

The group C structures (C1 to C3, 30% of A-D particles) show a change in the direction of SNF2H movement, compared to group B structures. SNF2H rotates downward and outward, away from the histone octamer, while the brace helix continues its outward movement initiated in the group B structures (**Figure 3a**). In group C structures, SNF2H adopts a canonical open conformation observed in group B and in structures of chromatin remodelers bound to ADP ^11^, and moves as a rigid body relative to the nucleosome (**Figure S7a**). These movements pull the guide strand of DNA outward at SHL3 and correct the distortion at SHL2 and SHL3 seen in group B structures (**Figure 3b**). Compared to the canonical nucleosome structure, both DNA strands in group C are displaced outward by more than 6 Å, taking a longer path that accommodates one additional base pair at SHL2 (**Figure 3c**). Thus, the group C structures show that the nucleosomal DNA is effectively moved by 1 bp from entry site to SHL2 (**Figure S7b**).

**Figure 3.**
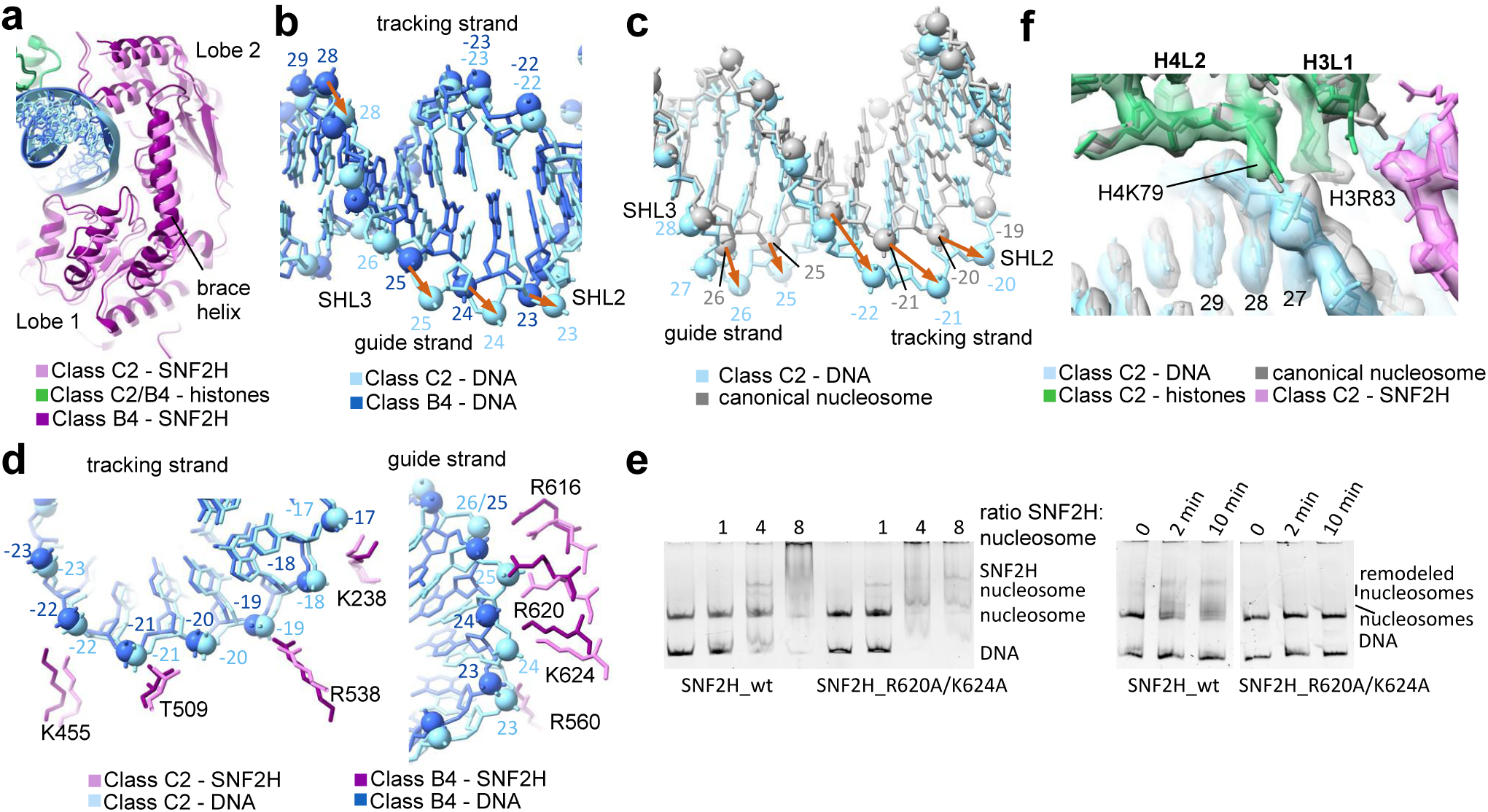
Tracking strand translocation. **a, b)** Close-up views of the overlay of class C2 (light pink) and B4 (purple, dark blue) structures, showing conformational changes in SNF2H (a) or in the DNA (b). In panel b, the pulling out of guide strand nucleotides is indicated with orange arrows. **c)** Overlay of the DNA at SHL2 and SHL3 in class C2 (light blue) and in the canonical nucleosome (grey) structures. The pulling out of the nucleosomal DNA is indicated with orange arrows. **d)** Close-up views of the overlay of class C2 (light pink, light blue) and B4 (purple, dark blue) structures, showing changes in SNF2H interaction with DNA tracking (left) and guide (right) strands. **e)** Native gel stained for DNA showing nucleosome binding (left) and remodeling (right) by wild-type and R620A K624A mutant SNF2H. **f)** Close-up view of the overlay of class C2 (color) and canonical nucleosome (grey) structures, showing changes in histones interacting with DNA at SHL2 and SHL3.

The three group C structures show gradual movements of SNF2H (**Figure S7c-e**). Despite those movements, the interactions between SNF2H and DNA nucleotides remain unaltered between groups B and C (**Figure 3d**). The movement of the brace helix away from the nucleosome (**Figure 3a and S7c-f**) pulls the guide strand at SHL3 DNA further outward (**Figure 3b, d and S7g-i**). The guide strand completes the motion initiated in group B and moves by 1 nt relative to canonical nucleosome (**Figure S7b, j**). This movement corrects the DNA distortion seen in group B, allowing the DNA to adopt the canonical conformation (**Figure 3c**). We mutated brace helix residues R620 and K624 to Ala and observed severe reduction in chromatin remodeling activity of SNF2H (**Figure 3e**), whereas the mutations had only a small impact on the ATPase (**Figure S7k**) or nucleosome binding activity (**Figure 3e**). We obtained similar results when we mutated brace helix residue R616 (**Figure S7k**). Together, these data support the essential role of these brace helix residues in DNA translocation.

These DNA rearrangements relieve most of the histone distortions seen in group B structures. For example, the nucleotide in position 28 completes its movement to position 27 in C2. Consequently, H3R83, which interacted with DNA at position 28 in group B structures, moves back to its canonical conformation in C2 and interacts with DNA at position 27 (**Figure 3f**). Still, some histone distortions remain in H3 L1 and H4 in group C structures (**Figure 3f and S7m**), since the DNA is still pulled away from the histone octamer surface compared to the canonical nucleosome.

### DNA release

Our structural analyses captured the initial step of DNA release from SNF2H in the single group D structure (10% of A-D particles). Compared to the group C structures, group D shows both lobes of SNF2H continuing their outward movement away from SHL3 and toward SHL2 (**Figure 4a and S8a-b**). Moreover, lobe 2 moves upward and away from lobe 1, opening the two lobes further apart (**Figure S8c**). Notably, we can still observe density consistent with ADP between the two lobes (**Figure S3a**). The brace helix continues its outward movement, away from the nucleosome; compared to A1 structure, it moves by more than 10 Å (**Figure 4a and S8d**). This movement of the brace helix weakens its interaction with the guide strand, allowing the DNA to move 1–2 Å back toward the surface of the histone octamer and the dyad (**Figure 4b-c and S8e-g**). Together, these observations indicate that opening of two lobes and brace helix movement lead to DNA release from SNF2H.

**Figure 4.**
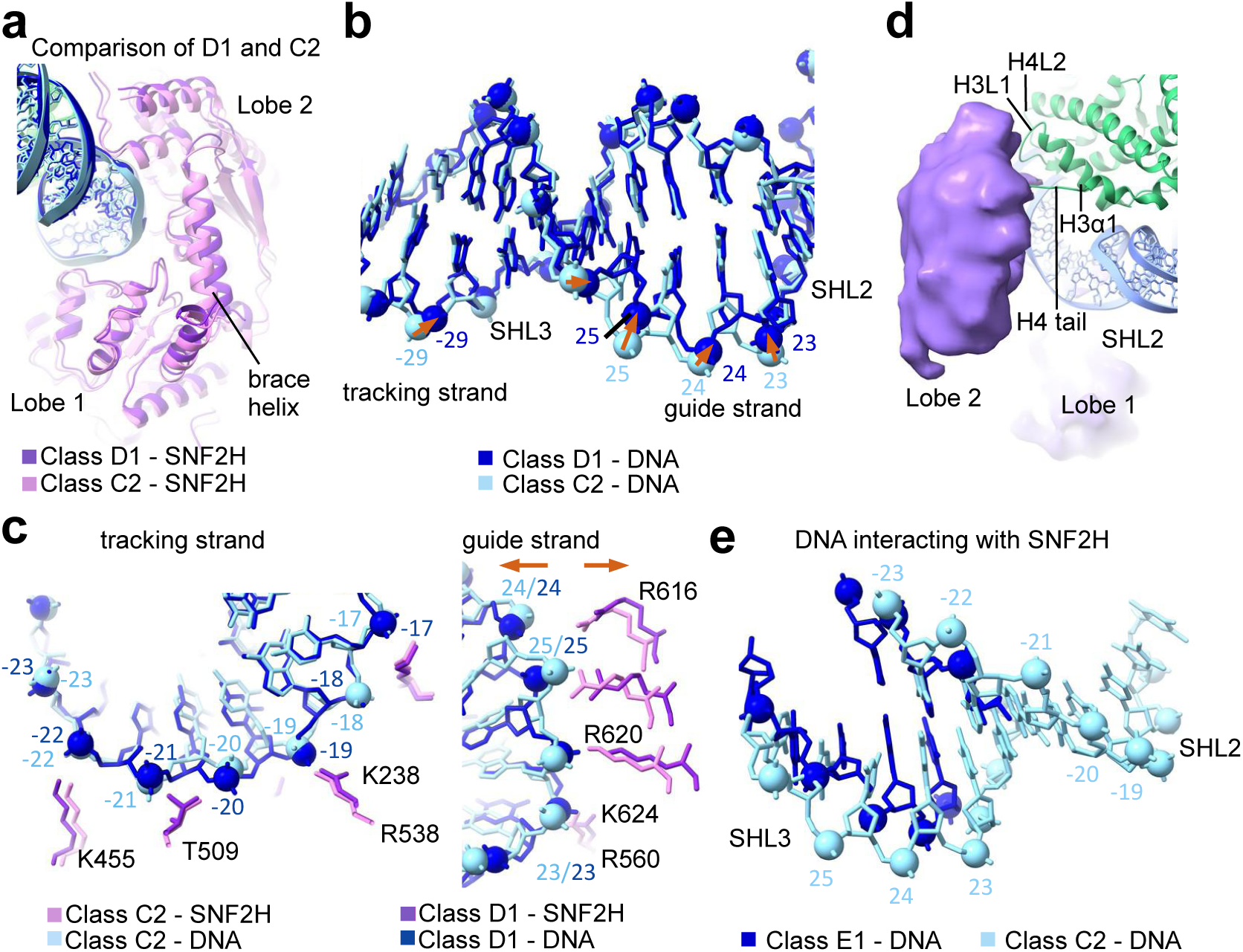
DNA release. **a)** Close-up view of the overlay of class D1 (purple) and C2 (light pink) structures, showing conformational changes in SNF2H. **b)** Close-up view of the overlay of the DNA at SHL2 and SHL3 in class D1 (dark blue) and C2 structures (light blue). Movement of nucleotides in the guide strand toward the histone core surface is indicated with orange arrows. **c)** Close-up views of the overlay of class D1 (purple, dark blue) and C2 (light pink, light blue) structures, showing interactions between SNF2H and DNA. **d)** Close-up view of class E structure showing interactions of SNF2H (purple) with DNA (blue) and histones (green). **e)** Overlay of the DNA at SHL2 and SHL3 in class E1 (dark blue) and C2 (light blue) structures, showing only the nucleotides that interact with SNF2H in each class.

In the two group E structures, the two lobes of SNF2H are fully separated from each other, but each lobe is still flexibly bound to the nucleosome (**Figure 1a, 4d and S8h**). Though the resolution for SNF2H is limited in these structures, the separation of the two lobes suggests that the nucleotide has dissociated from SNF2H. Lobe 2 continues its upward movement toward histones observed in the group D structure and releases the DNA at SHL2, which returns to the conformation in the canonical nucleosome (**Figure 1a, 4e**). Notably, SNF2H lobe 2 maintains its interactions with histones H3 and H4 (**Figure 4d**) and with DNA near SHL3 in the E1 structure (**Figure 4e and S8i-j**).

## Discussion

Our 13 structures illustrate the continuous motions of DNA, histones and SNF2H during chromatin remodeling. They show that conformational changes in SNF2H induced by ATP hydrolysis pull the nucleosomal DNA at SHL2 in two distinct steps (from A to B structures and from B to C structures), resulting in the formation of a DNA bulge that accommodates one additional base pair at SHL2. Subsequently, we observe opening of the SNF2H lobes and partial release of the DNA at SHL2 (C to D structures), followed by full separation of the two lobes and DNA release (group E) (**Figure S9a**). While each transition involves subtle conformational changes in SNF2H, they contribute to substantial movements: most of the lobe 2 and parts of lobe 1 move by 5 Å or more between A1 and D1 structures; the brace helix moves by more than 10 Å between A1 and D1 structures. Notably, the SNF2H residues that contact DNA at SHL2 during these translocation steps (as observed in our structures and validated by mutagenesis) are conserved across different families of chromatin remodelers, suggesting that the mechanisms we describe for SNF2H likely apply to all ATP-dependent chromatin remodeling enzymes (**Figure S9b**).

Previous structures had captured Snf2 and ISWI chromatin remodelers bound to non-hydrolyzable ATP analog ADP-BeFx ^11,12,15^. Our group A structures show overall similarity to the ADP-BeFx bound structures, but we observe gradual rotation of lobe 2, suggesting that our structures represent states that occur during ATP hydrolysis or before Pi release. In the group B structures SNF2H shows overall similarity to structures of Snf2 or ISWI bound to ADP ^11,12^, but it adopts a different position relative to the nucleosome. The ADP-bound structures of ISWI and Snf2 showed asynchronous translocation of the tracking strand by 1 nucleotide from entry site to SHL2. In contrast, our group B structures show both DNA strands translocate together from entry site until SHL3, with asynchronous translocation occurring only at SHL2 and SHL3, where movement of guide strand is blocked by the brace helix. These differences suggest that the ADP-bound state described previously ^11,12^ might be a short-lived intermediate that exists between our groups A and B structures but that we did not capture with actively translocating SNF2H. DNA distortion at SHL2 and SHL3 and formation of short A-DNA helix at SHL2 were previously observed in the structure of a nucleosome with chromatin remodeler Chd1 in a nucleotide-free state ^7^. In agreement, our structures show a short A-DNA helix at SHL2 in group B structures, as reported by the modeling algorithm 3DNA ^22^.

Our group B structures also show that histones adjust to the DNA movements induced by SNF2H activity, which reconciles conflicting observations in the literature. A previous report showed that cysteine substitutions and cross-linking of residues in H3 L1 (H3L82C) and H4 L2 (H4V81C) reduced SNF2H and ISW2 activity, indicating that structural alterations in histones are important for chromatin remodeling ^17,23^. However, histone conformational changes had not been observed in previous structures of chromatin remodelers stabilized by different nucleotides ^18,19^. The intrinsic plasticity of histones was previously described: we reported spontaneous histone deformation around SHL2 in nucleosomes without any remodelers ^24,25^; dynamic behavior of histones has been observed in recent MD simulations ^26^. Our cryo-EM structures of SNF2H actively remodeling the nucleosome show that residues in histones H3 and H4 undergo conformational changes (both at backbone and side chain level) in order to maintain their interactions with moving DNA phosphates. The histone backbone changes we observe are relatively small (1-2 Å) and although both residues H3L82 and H4V81 rearrange in B4 structure, the distance between them remains similar to the canonical nucleosome structure, suggesting that cross-linking those residues might underestimate functional importance of conformational changes in histones ^18,19^. However, larger structural changes may occur more transiently, and it is possible that cross-linking those positions might disrupt larger motions of H3 L1 and H4 L2 that occur during DNA translocation.

Previous Förster resonance energy transfer (FRET) experiments have shown an ATP-dependent delay between DNA entry into and exit from the nucleosome during remodeler-induced translocation ^27^. This is supported by our cryo-EM observations of actively translocating SNF2H: the majority of our particles (75%) belong to groups B and C, which indicates that, under our reaction conditions, the enzyme spends most of the time in those states. Thus, our observations are consistent with a delay between DNA movement at the entry site (class B1) and translocation toward the exit site (after class D1).

Here we interpret the group E structures as states during the DNA release step, but we fully acknowledge that alternative interpretations can be entertained. For example, group E structures could correspond to the initial association of SNF2H with the nucleosome, its dissociation from the nucleosome, or a regulatory state that does not occur at every DNA translocation cycle. Notably, in the E1 structure, SNF2H interacts with histones and DNA near SHL3, and these contacts with the nucleosome could provide directionality to the DNA movement, as they would prevent propagation of the bulge toward the entry site when the SHL2 DNA is released — thus, the additional 1 bp accumulated at SHL2 can only propagate in the opposite direction, toward the dyad.

## Methods

### Histone expression and purification

All the *Xenopus laevis* histones were separately over-expressed in BL21(DE3) pLysS bacterial strain and purified from inclusion bodies, as previously described ^28^.

The transformed cells were grown at 37 °C and induced with 1 mM IPTG at OD_600_ of 0.6. After 3h of induction, the cells were harvested, resuspended in lysis buffer (50 mM Tris-HCl (pH-7.5), 150 mM NaCl, 1 mM EDTA, 1 mM DTT and 0.1 mM PMSF) and frozen. The frozen cells were thawed and sonicated. The pellet containing inclusion bodies was recovered by centrifugation at 35,000 rcf for 20 min at 4 °C. The recovered pellet was washed two times with lysis buffer containing 1% Triton X-100 followed by two times wash with lysis buffer without Triton X-100. The pellet, containing histones as insoluble inclusion bodies was retrieved by centrifugation after each washing step.

The inclusion body pellet for each histone protein was extracted in a buffer containing 50 mM Tris (pH 7.5), 2 M NaCl, 6 M guanidine hydrochloride and 1 mM DTT for overnight at room temperature. Any insoluble components were removed by centrifugation. Histone pairs (H2A-H2B and H3-H4) were combined in equimolar ratios and dialyzed twice in 1L of refolding buffer (25 mM HEPES/NaOH (pH 7.5), 2 M NaCl and 1 mM DTT) at 4 °C. Precipitates were removed by centrifugation for 20 minutes at 13,000 rpm at 4 °C. The soluble histone pairs were purified by cation-exchange chromatography in batch (SP Sepharose Fast Flow resin). The samples were diluted fourfold with buffer without salt (25 mM HEPES/NaOH (pH 7.5) and 1 mM DTT) and bound to the resin for 30 min. The resin was extensively washed with 500 mM salt buffer in batch (25 mM HEPES/NaOH (pH 7.5), 500 mM NaCl and 1 mM DTT) and loaded onto a disposable column. On the column, the resin was washed, and pure proteins were eluted with 25 mM HEPES/NaOH (pH 7.5), 2 M NaCl and 1 mM DTT. Soluble histone pairs were concentrated and purified on a Superdex S200 size-exclusion column (GE) equilibrated in 25 mM HEPES/NaOH (pH 7.5), 2 M NaCl and 1 mM DTT. Clean protein fractions were pooled, concentrated and flash frozen.

### Histone octamer preparation

Histone octamer purification was done using the following protocol ^28,29^. In short, a 2-fold molar excess of the H2A-H2B dimer was mixed with the H3-H4 tetramer in the presence of buffer IV containing 2 M NaCl, 25 mM HEPES (pH 7.5) and 1 mM DTT. After overnight incubation at 4 °C, octamer was separated from excess dimer using a Superdex S200 Increase 10/300 GL column on AKTA FPLC equilibrated with buffer IV. The fractions of interest were analyzed on SDS-PAGE, pooled and concentrated for final nucleosome assembly.

### Nucleosomal DNA preparation

Nuclesomal DNA was prepared by PCR amplification from a plasmid with 145 base pairs of 601 DNA sequeunce ^30^. Primers were designed such that the final DNA product had a 80 base pair long overhang on one side of 601 DNA. The sequence of 601 with 80 bp of flanking DNA is as following: GCACAGGATGTATATATCTGACACGTGCCTGGAGACTAGGGAGTAATCCCCTTGGCGGT- TAAAACGCGGGGGACAGCGCGTACGTGCGTTTAAGCGGTGCTAGAGCTGTCTACGAC- CAATTGAGCGGCCTCGGCACCGGGATTCTCCAGGGCGGCCGCGTATAGGGTCCATCA- CATAAGGGATGAACTCGGTGTGAAGAATCATGCTTTCCTTGGTCATTAGGATCC

### Nucleosome assembly

Nucleosome assembly was done by double bag dialysis as previously described ^31^. Nucleosomal DNA was suspended in 25 mM HEPES/NaOH pH 7.5, 1 mM DTT, 2 M NaCl. The histone octamer was mixed with DNA in equimolar ratios and placed into a dialysis button (membrane cutoff: 3.5 kDa MW). These buttons were then placed in a dialysis bag (6-8 kDa cut off membrane) filled with 50 ml of buffer containing 25 mM HEPES (pH 7.5), 2 M NaCl and 1 mM DTT. The dialysis bag was immersed into 1 l of buffer containing 25 mM HEPES (pH 7.5), 1 M NaCl and 1 mM DTT and dialyzed overnight at 4 °C. After overnight dialysis, the 1L buffer is replaced with 1L of 25 mM HEPES (pH 7.5), 50 mM NaCl, and 1 mM DTT for a 6 hours dialysis. Finally, the last dialysis is done with 25 mM HEPES (pH 7.5) and 1mM DTT for an hour. The quality of nucleosome assemblies was checked on a 6% native PAGE which is run on 1x TBE, 200V at 4 °C. The gels were visualized on a Chemidoc.

### Purification of SNF2H

SNF2H cloned in pET duet vector was ordered from GenScript. *E. coli* LOBSTR Rosetta bacterial culture, containing pET duet vector with SUMO tag on one site followed by SNF2H gene, was grown in TB medium containing the appropriate antibiotics at 37 °C until the optical density reached 0.6 at 600 nm. The culture was induced with 0.4 mM IPTG for overnight at 16 °C. The cells were harvested and frozen at −20 °C. For purification, frozen pellets were thawed and resuspended in lysis buffer (20 mM Tris-HCl/ HEPES pH 8.0, 2 M NaCl, 1 mM PMSF, 10 mM Imidazole, 1 mM DTT) and lysed. The cleared supernatant was incubated with pre-equilibrated Ni Sepharose 6 Fast flow resins. On the column, the resin was washed with 5 bed volumes of washing buffer I (20 mM Tris-HCl/HEPES pH 8.0, 2 M NaCl, 10mM Imidazole 1 mM DTT), II (20 mM Tris-HCl pH 8.0, 150 mM NaCl, 75/50 mM Imidazole, 1 mM DTT) and III (20 mM Tris-HCl/HEPES pH 8.0, 150 mM NaCl, 300/500 mM Imidazole, 1 mM DTT). The bound protein was eluted with 5 bed volumes of buffer III and 5 column volume of 1M imidazole. The Ni purified protein was then purified on SP Sepharose cation exchange resins using 20 mM Tris-HCl/HEPES pH 8.0, 1 M NaCl and 1 mM DTT as the elution buffer. The protein was further treated with PreScission Protease, followed by overnight dialysis with buffer containing 20 mM HEPES pH 7.5, 300mM NaCl, 1 mM DTT. The protein was further concentrated and applied on size exclusion Superdex 200 Increase 10/300 equilibrated in 20 mM HEPES/NaOH, pH 7.5, 150 mM NaCl, 1 mM DTT. The fractions with the protein were analyzed on SDS-PAGE and concentrated. The concentrated protein was flash frozen and stored at −80 °C. SNF2H mutant variants were created using site directed mutagenesis and purified using same protocol as described above for SNF2H(WT). Quantification of SNF2H variants were performed using SDS-PAGE for assays.

### Nucleosome Cross-linking

Nucleosome cross-linking was done by incubating assembled nucleosomes with 0.1% glutaraldehyde for 6 minutes at room temperature ^24,25^ and quenching with 50 mM Tris/HCl pH 7. The cross-linked nucleosome was analyzed on SDS-PAGE gel.

### Salt disassembly of the nucleosome

To test if DNA was cross-linked to histones, 7 μl of nucleosome sample (25 mM HEPES/NaOH pH 7.5, 1 mM DTT) was supplemented with the buffer containing NaCl to achieve different NaCl concentrations (0, 0.1, 0.2, 0.3, 0.5, 0.7, 1, and 1.5 M) ^24,25^. These samples were then incubated at 25 °C for 30 minutes. 4% glycerol (final concentration) was added to the samples and analyzed using 5% native PAGE. The gel was stained with SYBR Gold.

### SNF2H binding to nucleosome

The binding assay for SNF2H and nucleosome was performed at room temperature in 20 mM Tris (pH 7.5), 70 mM KCl, 1mM AMPPNP, 5 mM MgCl_2_ and 0.02% NP-40. Typically, 50 nM nucleosome (601+80) was incubated with different SNF2H at different concentrations (50 nM, 250 nM, 500 nM and 1 mM) as indicated and incubated for 40 minutes. The bound and unbound nucleosomes were separated on a 5% native polyacrylamide gel and stained with SYBr gold dye. The gels were imaged using Chemidoc.

To analyze the binding of SNF2H wild-type and mutants to the nucleosome, 4nM nucleosome and different concentrations of SNF2H (4 nM, 16 nM, and 32 nM) were used.

### SNF2H-remodeling assay

Nucleosome remodeling was done with 30 nM nucleosome and 60nM (two-fold molar excess) of SNF2H. The assay was carried out at room temperature in 20 mM Tris (pH 7.5), 70 mM KCl, 1mM ATP, 5 mM MgCl_2_ and 0.02% NP-40. This assay was also carried out in presence of 0.25 mM or 0.5 mM MgCl_2_ to check the remodeling activity under reduced Mg^2+^ concentrations. The reaction was stopped with excess of ADP/AMP-PNP and 50 mM EDTA. The reaction products were loaded on 3-12% bis-tris acrylamide gels and the gels were visualized after SYBr gold staining on a Chemidoc.

This method of visualization was followed for all the remodeling assays unless stated otherwise. To analyze the remodeling activity of SNF2H mutants, 4 nM of nucleosome and 4 nM of wt or mutant SNF2H were used. The assay was done as described above and the reaction products were analyzed on 5% native polyacrylamide gels. To check the remodeling of cross-linked nucleosomes, 4 nM of nucleosomes and 4 nM of SNF2H were used and reaction products were run on a 5% native polyacrylamide gel and visualized as described above.

### ATPase assay

To measure the ATPase activity of SNF2H, ADP-glo^TM^ kinase assay kit from Promega was used. This kit measures the amount of ADP released by an ATPase. 32 nM of SNF2h wild type or mutant was incubated with 4 nM of nucleosome at room temperature for 20 minutes, with the same buffer used in remodeling assays. Two-fold serial dilutions of the complex were incubated with 1 mM ATP. The signal was measured at 590 nm using POLstar omega from BMG Labtech.

### Assembly of SNF2H-nucleosome complex for cryo-EM analysis and grid preparation

Nucleosomes, with the final concentration not exceeding 0.3 mM, were assembled by ‘double bag’ dialysis using the same steps as described in the “Nucleosome assembly” section. 2.5 μM SNF2H and 0.625 μM nucleosomes were mixed in such a way that the final salt concentration did not exceed 50 mM NaCl. The binding was done for 20 minutes at room temperature and monitored as described in the binding assay section except for Mg^2+^ concentrations. 3 ml of SNF2h-nucleosome complex was applied to freshly glow-discharged Quantifoil R1.2/1.3 holey carbon grid. For freezing, FEI Vitrobot Mark IV chamber was maintained at 95% humidity and temperature of +12 °C. After 3 s blotting time, grids were plunge-frozen in liquid ethane using FEI Vitrobot automatic plunge freezer. SNF2H was frozen at 5’’, 2’ and 10’ after addition of ATP at two different Mg^2+^ concentrations, 0.25 mM and 0.5 mM. For the 5’’ time point, 1 mM ATP was added to the sample that was already on the grid and the sample was immediately frozen. For 2’ and 10’ timepoints, 1mM ATP was added prior applying to the grid.

### Cryo-EM data collection, image processing and model building

Electron micrographs were recorded on FEI Titan Krios at 300 kV with a Gatan Summit K3 electron detector using EPU at the cryo-EM facility at St. Jude Children’s Research Hospital. Image pixel size was 0.6485 Å per pixel on the object scale. Data were collected in a defocus range of 14,000 – 30,000 Å with a total exposure of 60 e^−^ Å^−2^. We collected ∼20 000 images of the sample frozen 5 seconds after activation (0.25 mM Mg^2+^), ∼23 000 and 48 000 of the sample frozen 2’ after activation (0.25 mM and 0.5 mM Mg^2+^ respectively) and ∼20 000 frames 10’ after activation. Micrographs were aligned with the MotionCorr2 software using a dose filter ^32,33^. The contrast transfer function parameters were determined using CTFFIND4 ^34^.

Particles were picked using TOPAZ in RELION software package ^35,36^. Particles were binned and 2D class averages were generated in RELION. Inconsistent class averages were removed from further data analysis. The initial reference was filtered to 40 Å in RELION. C1 symmetry was applied during refinements for all classes. Particles were split into many datasets and refined independently, and the resolution was determined using the 0.143 cut-off (RELION auto-refine option). All maps were filtered to resolution using RELION with a B-factor determined by RELION. Particles containing SNF2H (class 2) were classified many times leading to 11 different structures that have overlapping particles due to continuous motion. Final maps were processed using EMReady ^37^.

Initial molecular models were built using Modelangelo and Coot ^38,39^. The model of the nucleosome (Protein Data Bank (PDB): 6WZ5) was refined into the cryo-EM map in PHENIX ^40,41^. The model of SNF2H bound to the nucleosome (PDB:6NE3) was either rigid body fitted using PHENIX or built into cryo-EM maps using Modelangelo, manually adjusted and rebuilt in Coot and refined in Phenix ^21^. Visualization of all cryo-EM maps were done in Chimera ^42^.

## Acknowledgments

We thank Cryo-EM Center members (Martin Turk, Sagar Chittori, Asfarul Haque, Jose Miguel de la Rosa Trevin) at St. Jude Children’s Research Hospital for support with grid screening and data collection. We thank Inês Chen for critical reading and comments, Zhaowen Luo for help with making movies, Ivica Zamarija for making DNA. Work in the Halic laboratory is funded by St. Jude Children’s Research Hospital, the American Lebanese Syrian Associated Charities, and NIH awards 1R01GM135599 and 1R01GM141694.

## Author contributions

M.H. designed the experiments. D.M. and A.D. purified wild type SNF2H. A.D. cloned and purified-SNF2H mutants. D.M. performed biochemical experiments. D.M. made grids and collected data with help from S.B. M.H analyzed cryo-EM data. D.M and A.D. built models. D.M, A.D, S.B. and M.H. analyzed the data. M.H. wrote the paper with help from all the authors.

## Competing interests

The authors declare no competing interests.

## Data availability

EM density maps and models have been deposited in the Electron Microscopy Data Bank and PDB under the following accession codes: for canonical nucleosome PDB 9E1Y was built using map EMD-47425, Class A1 EMD-47412 and PDB 9E1L, Class A2 EMD-47413 and PDB 9E1M, Class A3 EMD-47414 and PDB 9E1N, Class B1 EMD-47415 and PDB 9E1O, Class B2 EMD-47416 and PDB 9E1P, Class B3 EMD-47417 and PDB 9E1Q, Class B4 EMD-47418 and PDB 9E1R, Class C1 EMD-47421 and PDB 9E1U, Class C2 EMD-47422 and PDB 9E1V, Class C3 EMD-47423 and PDB 9E1W, Class D1 EMD-47424 and PDB 9E1X, Class E1 EMD-47427 and Class E2 EMD-47428. All other data supporting the findings of this study are available within the article and its supplementary information files.

---

**Figure S1.**
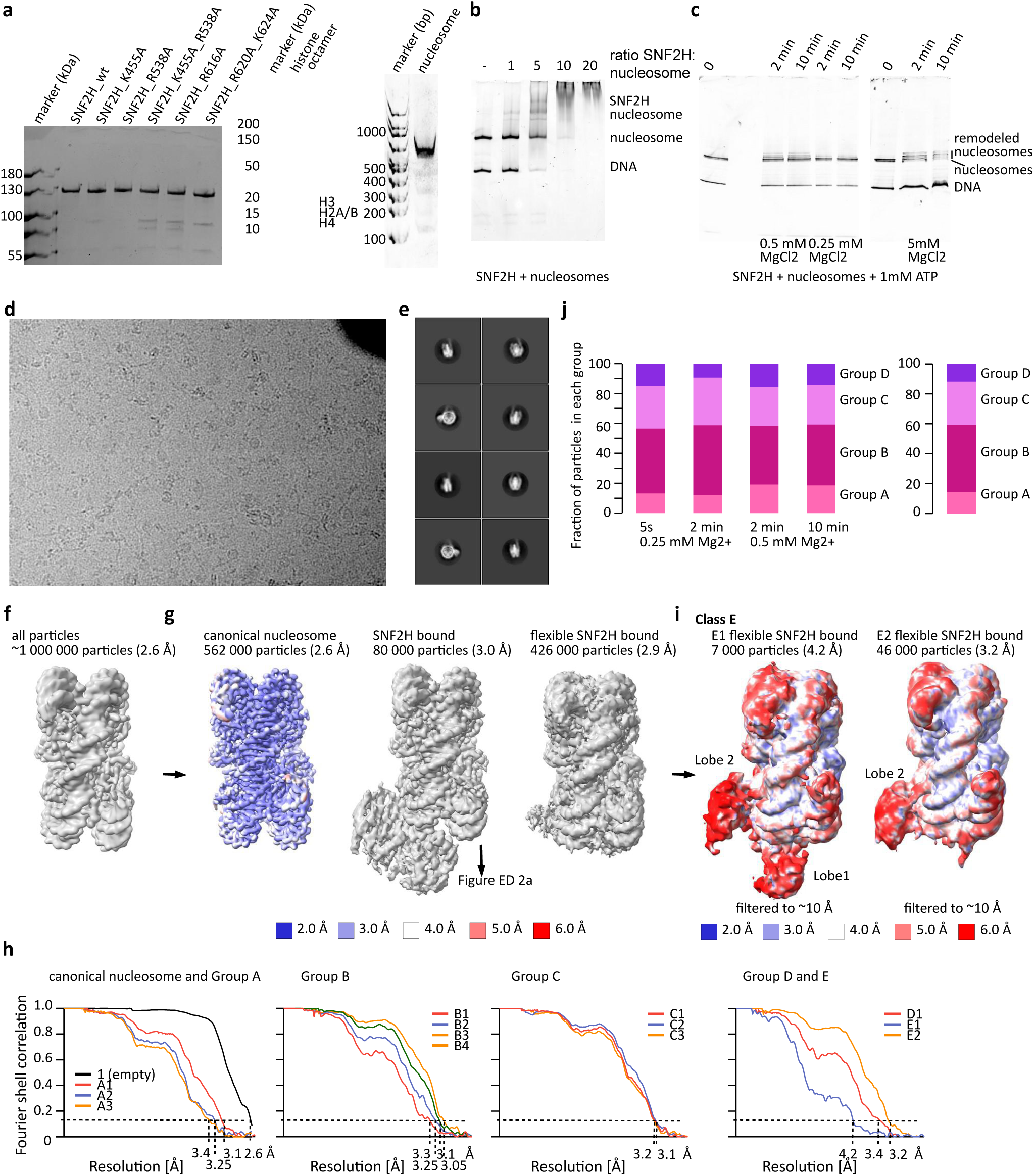
Assembly and EM analyses of SNF2H bound to nucleosome. **a)** SDS-PAGE showing purified human SNF2H (left, wild type and mutants) and assembled histone octamer assembly (middle). Right, native polyacrylamide gel stained for DNA shows the assembled nucleosome. **b)** Native polyacrylamide gel stained for DNA showing binding of SNF2H to the nucleosome. **c)** Native gel stained for DNA showing nucleosome remodeling by SNF2H upon addition of ATP. **d)** Representative cryo-EM micrograph from a set of ∼100,000 micrographs collected with Titan Krios electron microscope at 300 keV. Nucleosome particles in multiple orientations are visible. **e)** Representative 2D class averages showing nucleosomes with SNF2H bound in different orientations. **f)** Cryo-EM map of nucleosome from the entire dataset, refined to 2.6 Å. **g)** Classification of the data in panel 1f) resulted in three major classes of nucleosomes: nucleosome alone (canonical), SNF2H-bound nucleosome and flexible SNF2H-bound nucleosome. The number of particles and overall resolution of each class are shown. The map of the canonical nucleosome is colored by local resolution. **h)** Fourier shell correlation (FSC) curves showing the resolution of the maps shown in Fig S1g,i and S2a. **i)** Classification of flexible SNF2H-bound nucleosome data in panel g resulted in two classes, E1 and E2. The number of particles corresponding to each class and overall resolution are shown. Maps are colored by local resolution and filtered to 10 Å to visualize flexible SNF2H. **i)** Left, bar charts showing fraction of particles in each group (A to D) in the datasets collected at different time points and MgCl_2_ concentrations. Right, bar chart showing fraction of particles in each group in combined datasets (A-D).

**Figure S2.**
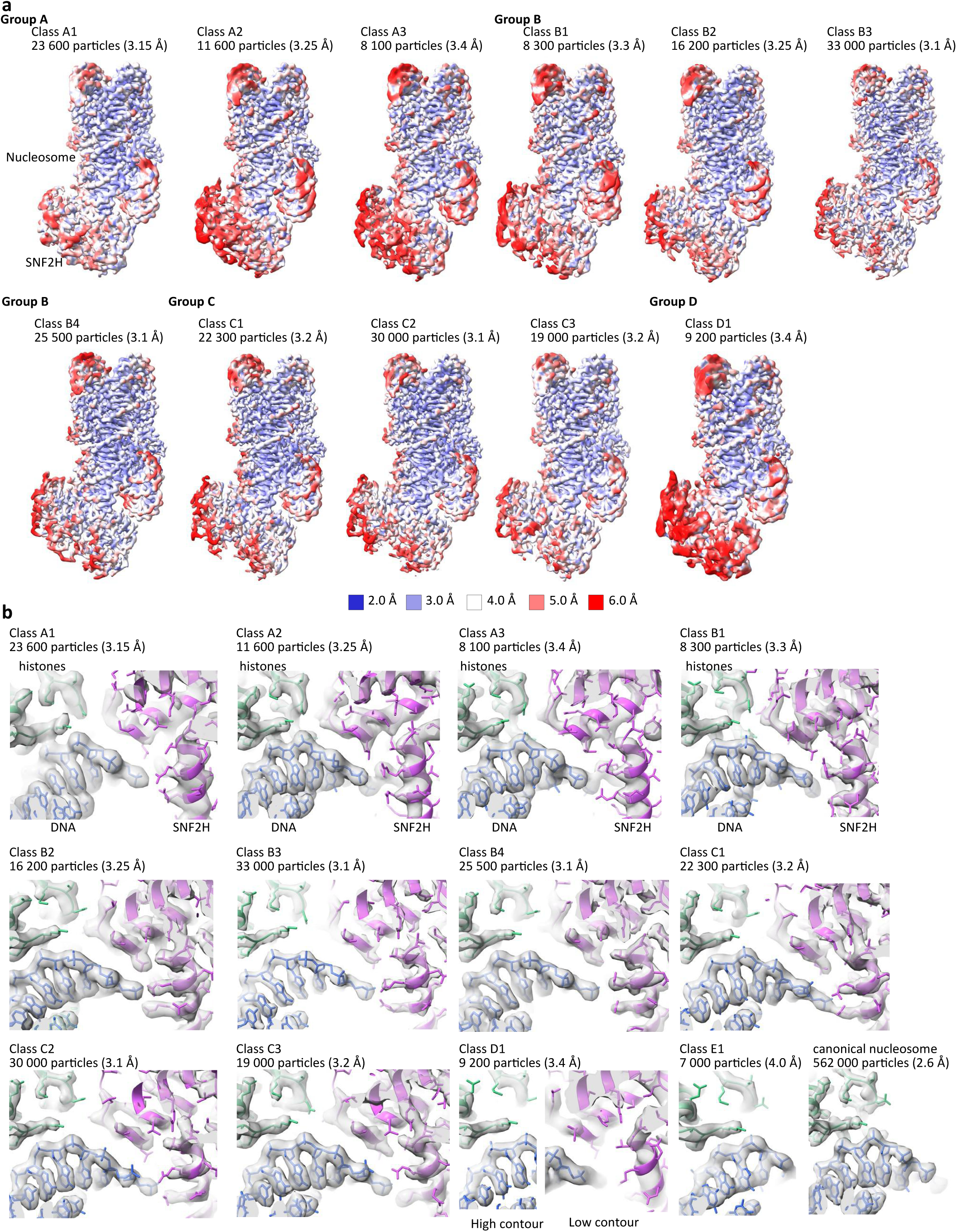
Classification of the SNF2H-nucleosome complex. **a)** Classification of SNF2H-bound nucleosome from ED 1g. Because of the continuous motion of SNF2H, many classifications were performed, resulting in 11 unique maps with overlapping particles. Maps are colored by local resolution. The number of particles and resolution in each class are shown. **b)** A representative region showing map quality and fit of the model is shown for the nucleosome and SNF2H from each map. DNA bases (blue) and histone side chains (green) are well resolved in all maps. Side chains in SNF2H (magenta) are resolved in most maps.

**Figure S3.**
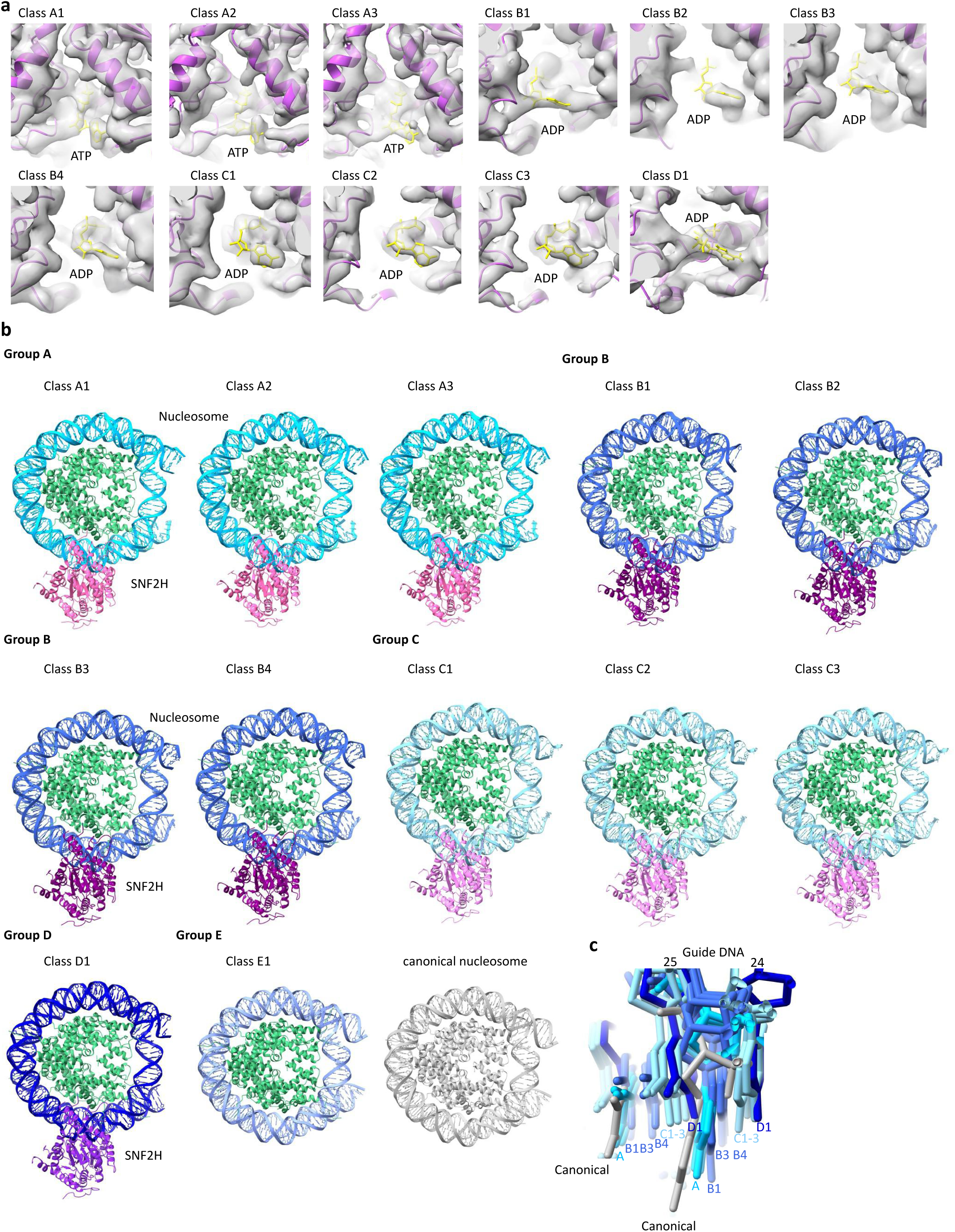
SNF2H bound to nucleosome. **a)** The nucleotide-binding site of SNF2H in all classes from groups A to D, showing map quality and fit of the model. Density for bound nucleotide is shown in yellow. **b)** Models of SNF2H bound to nucleosome are shown for all structures. Based on DNA conformation and SNF2H conformation, the 13 structures were grouped into 5 groups. **c)** Overlay of all models from groups A-D, showing the extent of the movement of nucleotides at positions 24 and 25 of the DNA guide strand.

**Figure S4.**
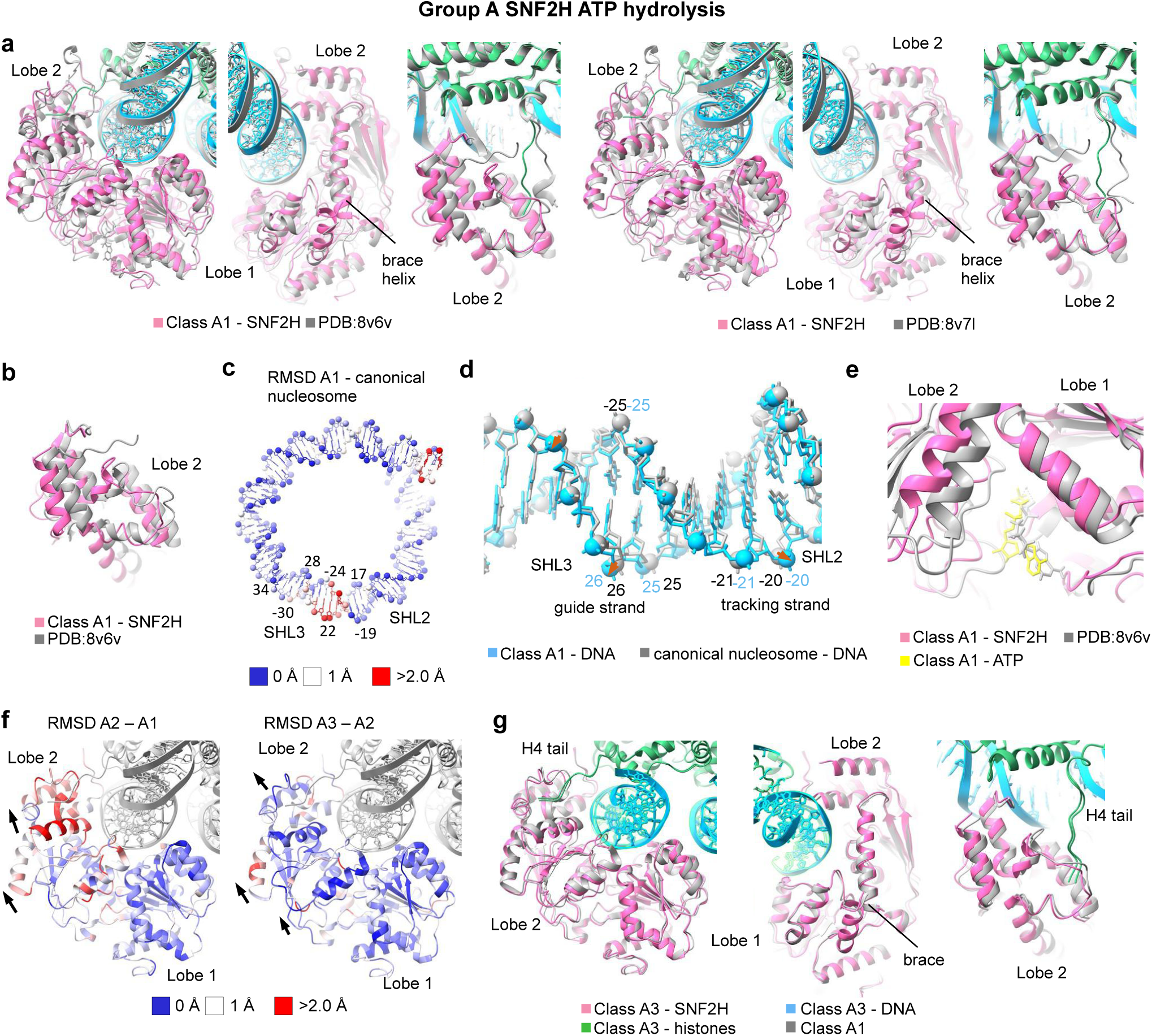
Conformational changes in SNF2H and nucleosome in group A. **a)** Close-up views of the overlay of class A1 (color) and two conformations of SNF2H–nucleosome complex with ADP-BeFx (grey; left, PDB 8v6v; right, PDB 8v7l). The structures were aligned on the nucleosome. **b)** Close-up views of the overlay of SNF2H lobe 2 from class A1 (color) and nucleosome-SNF2H with ADP-BeFx (grey, PDB 8v6v) structures. The structures were aligned on SNF2H lobe 2. **c)** RMSD of DNA between class A1 and canonical nucleosome structure, showing DNA changes upon SNF2H binding. Canonical nucleosomal DNA is shown. **d)** Overlay of the DNA at SHL2 and SHL3 in class A1 (bright blue) and canonical nucleosome (grey) structures. **e)** Close-up view of the overlay of SNF2H in class A1 (color) and ADP-BeFx-bound (grey, PDB:8v6v) structures, showing nucleotide-binding site. **f)** RMSD of SNF2H between class A2 and A1 (left, A1 shown) and A3 and A2 (right, A2 shown) structures. Black arrows show direction of the movement. **g)** Close-up views of the overlay of class A3 (color) and A1 (grey) structures, showing conformational changes in SNF2H.

**Figure S5.**
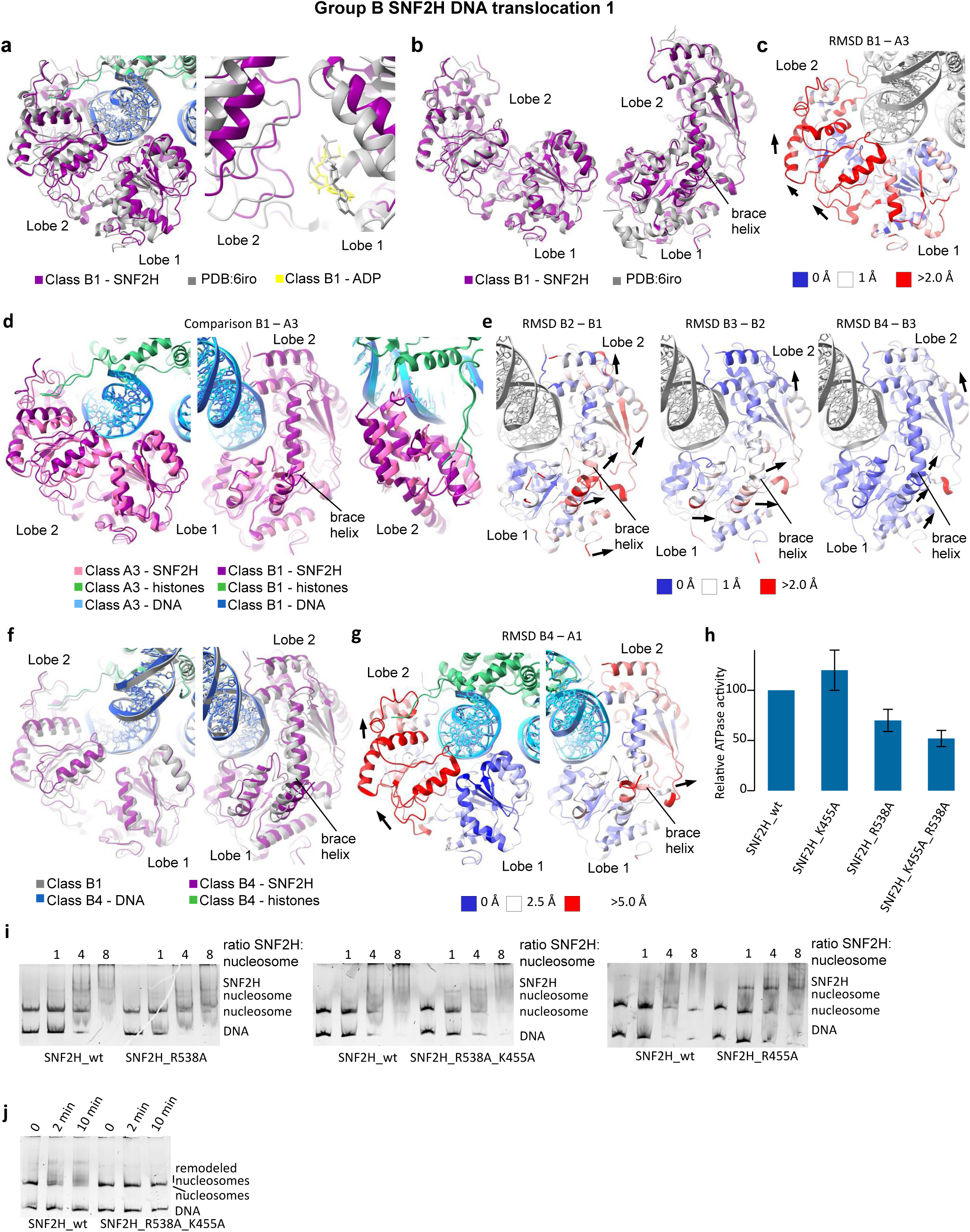
Conformational changes in SNF2H in group B. **a)** Close-up views of the overlay of class B1 (color) and nucleosome-SNF2H in ADP-bound state (grey, PDB:6iro) structures, showing conformational changes in SNF2H (left) and nucleotide binding site (right). Nucleosome was used for alignment. **b)** Close-up view of the overlay of SNF2H of class B1 (color) and SNF2H in ADP-bound state (grey, PDB:6iro) structures. SNF2H was used for alignment. **c)** RMSD of SNF2H between class B1 and A3 structures. A3 structure is shown. Black arrows show direction of the movement. **d)** Close-up views of the overlay of class B1 (purple, dark blue) and A3 (pink, light blue) structures, showing conformational changes in SNF2H. **e)** RMSD of SNF2H between class B2 and B1 (left, B1 shown) structures; B3 and B2 (middle, B2 shown); and B4 and B3 (right, B3 shown) structures. Black arrows show direction of the movement. **f)** Close-up views of the overlay of class B4 (color) and B1 (grey) structures, showing conformational changes in SNF2H. **g)** RMSD between the class B4 and A1 structures, showing rotation of SNF2H lobes 1 and 2 and brace helix. Class A1 is shown on both views; DNA and histone are in light blue and green, respectively. Black arrows show direction of the movement. **h)** ATPase activity of wild type and mutant SNF2H. ATPase activity of wild type is set to 1. **i)** Native polyacrylamide gel stained for DNA showing binding of SNF2H wild-type and mutants to the nucleosome. **j)** Native polyacrylamide gel stained for DNA showing nucleosome remodeling by SNF2H wild-type and double mutant R538A R455A.

**Figure S6.**
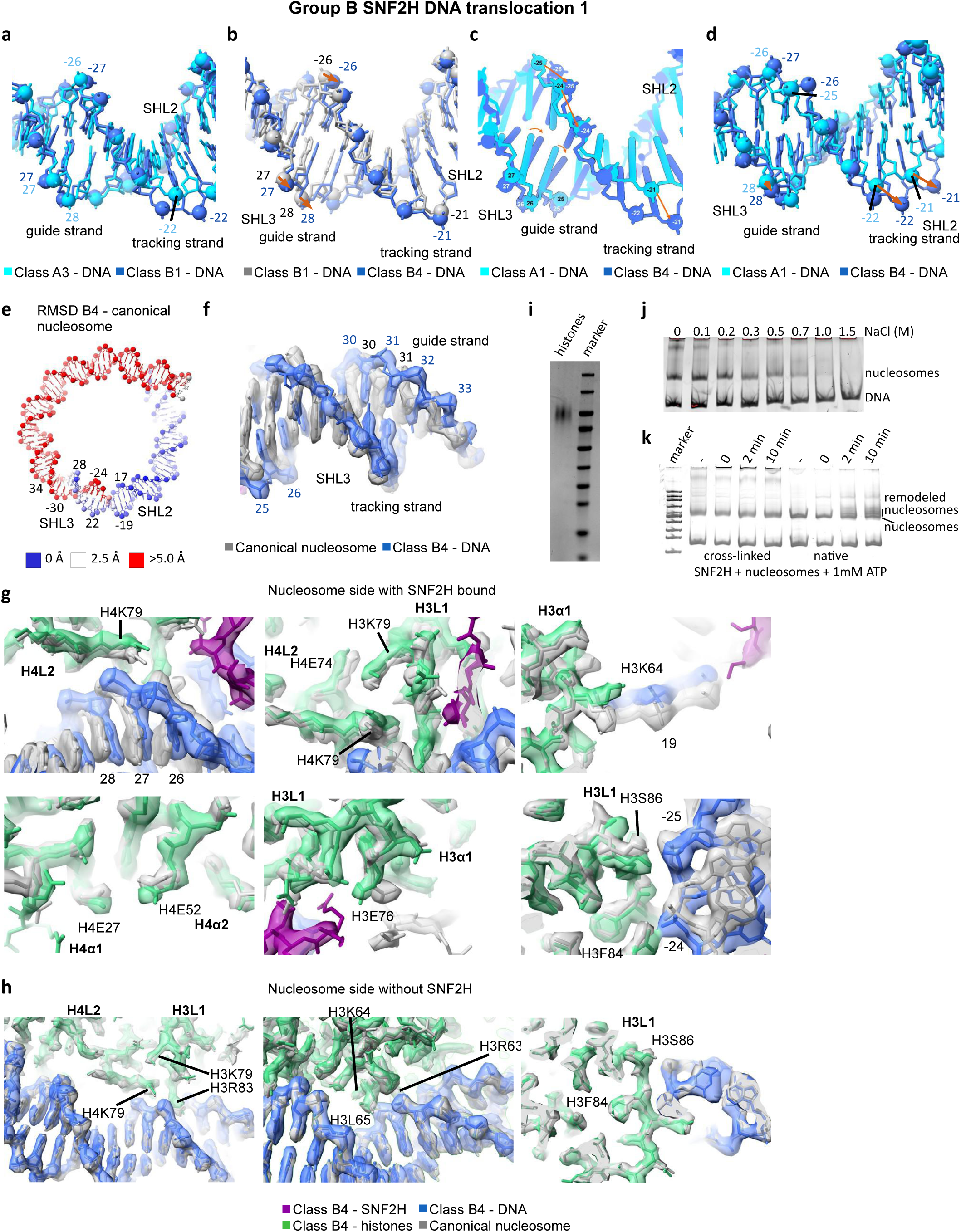
Conformational changes in nucleosome in group B. **a)** Overlay of the DNA at SHL2 and SHL3 in class B1 (blue) and class A3 (light blue) structures. **b)** Overlay of the DNA at SHL2 and SHL3 in class B4 (royal blue) and B1 (grey) structures. Orange arrows show direction of the movement. **c)** Cartoon showing overlay of the DNA at SHL2 and SHL3 in class B4 (royal blue) and A1 (light blue) structures. Same view as in Figure 2d. Orange arrows show direction of the movement. **d)** Overlay of the DNA at SHL2 and SHL3 in class B4 (royal blue) and A1 (light blue) structures. Orange arrows show direction of the movement. **e)** RMSD between the DNA in class B4 and class A1 structures. A1 DNA is shown. **f)** Overlay of the DNA at SHL2 and SHL3 in class B4 (royal blue) and canonical nucleosome structures (grey). **g, h**) Close-up views of the overlay of class B4 (color) and canonical nucleosome (grey) structures, showing DNA movement and resulting changes to histone residues interacting with DNA at SHL2 and SHL3 on SNF2H-bound side (f) and no changes to DNA or histones on the opposite side of the nucleosome (g). **i)** SDS-PAGE showing cross-linked histones migrating at 100 kDa. **j)** Native showing with salt-mediated DNA unwrapping from cross-linked nucleosome. These data show that DNA is not cross-linked to histones. **k)** Native gel stained for DNA showing remodeling of native (non cross-linked) and cross-linked nucleosomes by SNF2H.

**Figure S7.**
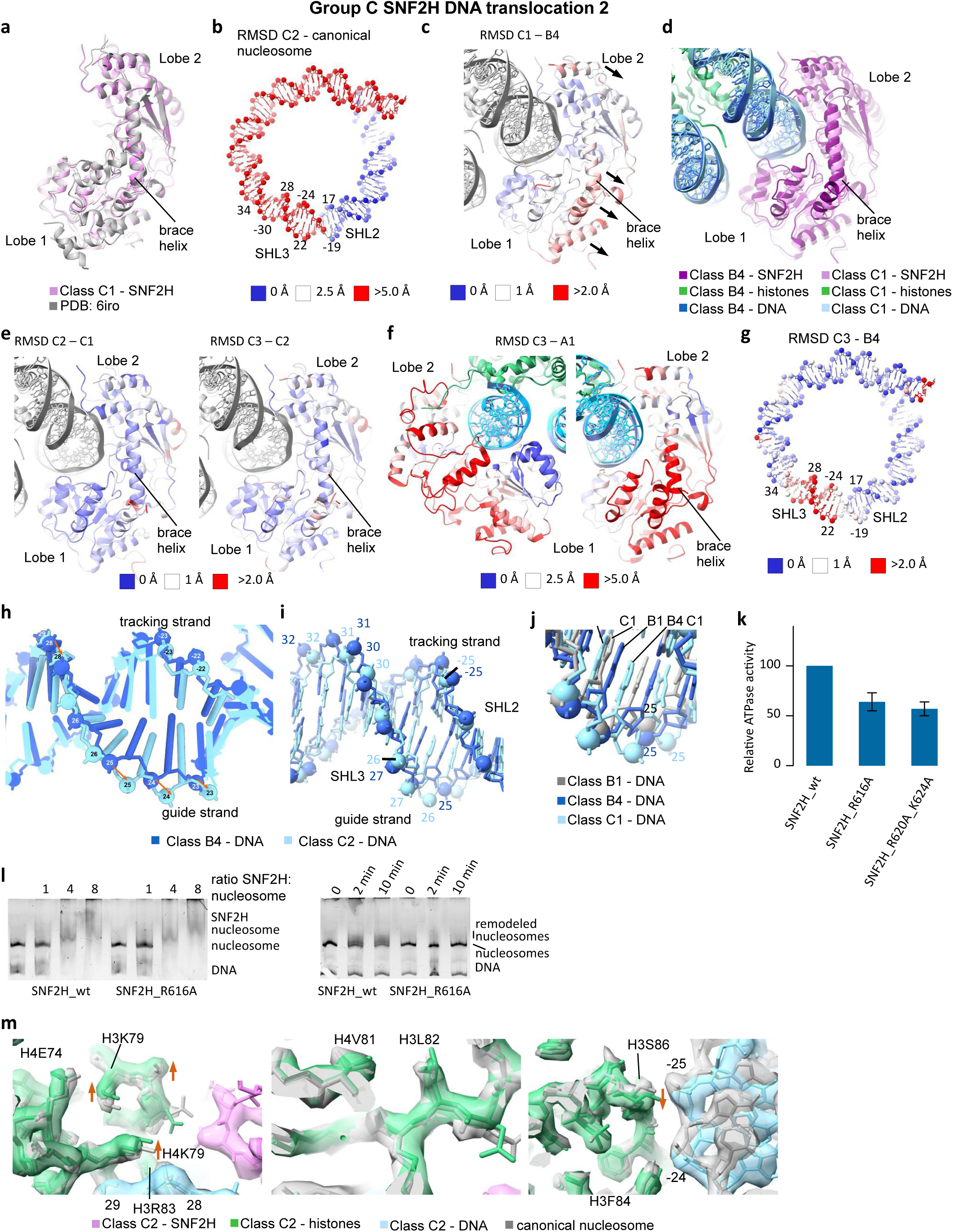
Conformational changes in SNF2H and nucleosome in group C. **a)** Overlay of SNF2H in class C1 (violet) and PDB:6iro (grey) structures. SNF2H was used for alignment. **b)** RMSD of DNA between class C2 and canonical nucleosome structures, showing DNA movement during translocation. Canonical nucleosome DNA is shown. **c)** RMSD of SNF2H between classes C1 and B4 structures. Class B4 is shown. Black arrows show direction of the movement. **d)** Close-up views of overlay of class C1 (light colors) and B4 (dark colors) structures, showing conformational changes in SNF2H (pink) and DNA (blue); histones shown in green. **e)** RMSD of SNF2H between classes C2 and C1 (left, C1 is shown) and C3 and C2 (right, C2 is shown) structures. **f)** RMSD of SNF2H between classes C3 and A1 structures, two views. Class A1 is shown. **g)** RMSD of DNA between class C3 and B4 (B4 is shown) structures, showing DNA movement in the second step of DNA translocation. **h)** Cartoon showing overlay of the DNA at SHL2 and SHL3 in class C3 (light blue) and B4 (royal blue). Same view as in Figure 3b. **i)** Overlay of the DNA at SHL2 and SHL3 in C2 (light blue) and B4 (royal blue) structures. **j)** Overlay of the DNA at SHL2 and SHL3 in C1 (light blue), B4 (royal blue) and B1 (grey) structures. **k)** ATPase activity of SNF2H wild type and mutants. ATPase activity of wild type is set to 1. **l)** Native polyacrylamide gel stained for DNA showing nucleosome remodeling (left) and binding (right) by SNF2H wild type and R616A mutant. **m)** Close up views of overlay of class C2 (color) and canonical nucleosome (grey) structures, showing DNA movements and resulting changes in histone residues interacting with DNA at SHL2 and SHL3. Orange arrows show direction of the movement.

**Figure S8.**
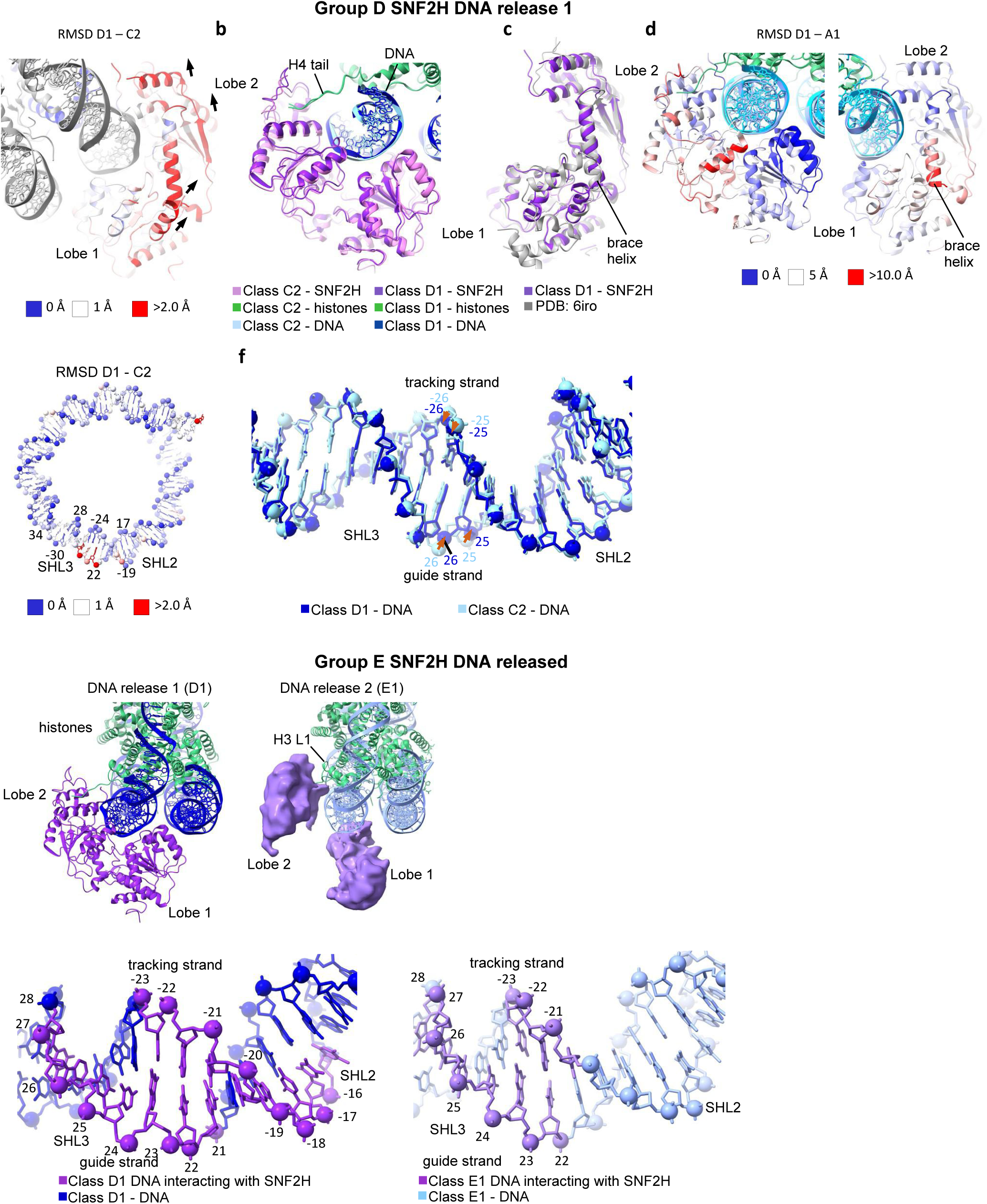
Conformational changes in SNF2H and nucleosome in groups D and E. **a)** RMSD of SNF2H between class D1 and C2 structures (C2 is shown). Black arrows show direction of the movement. **b)** Close-up view of overlay of class D1 (dark colors) and C2 (light colors) structures, showing conformational changes in SNF2H. Nucleosome was used for alignment. **c)** Close-up view of overlay of SNF2H in class D1 (dark violet) and PDB:6iro structures. SNF2H was used for alignment. **d)** RMSD of SNF2H between class D1 and A1 structures (A1 is shown). **e)** RMSD of DNA between class D1 and C2 structures, showing DNA movement in the first step of DNA release. DNA from C2 is shown. Orange arrows show direction of the movement. **f)** Overlay of the DNA at SHL2 and SHL3 at DNA release step 1 (D1, blue) and DNA translocation step 2 (C2, light blue). **g)** Close-up view of class D1 (left) and E1 (right) structures, showing separation of SNF2H lobes 1 and 2 in the latter. **h)** DNA at SHL2 and SHL3 in class D1 (left) and E1 (right) structures. For each structure, nucleotides that interact with SNF2H are colored purple; nucleotides that do not interact with SNF2H are colored blue.

**Figure S9.**
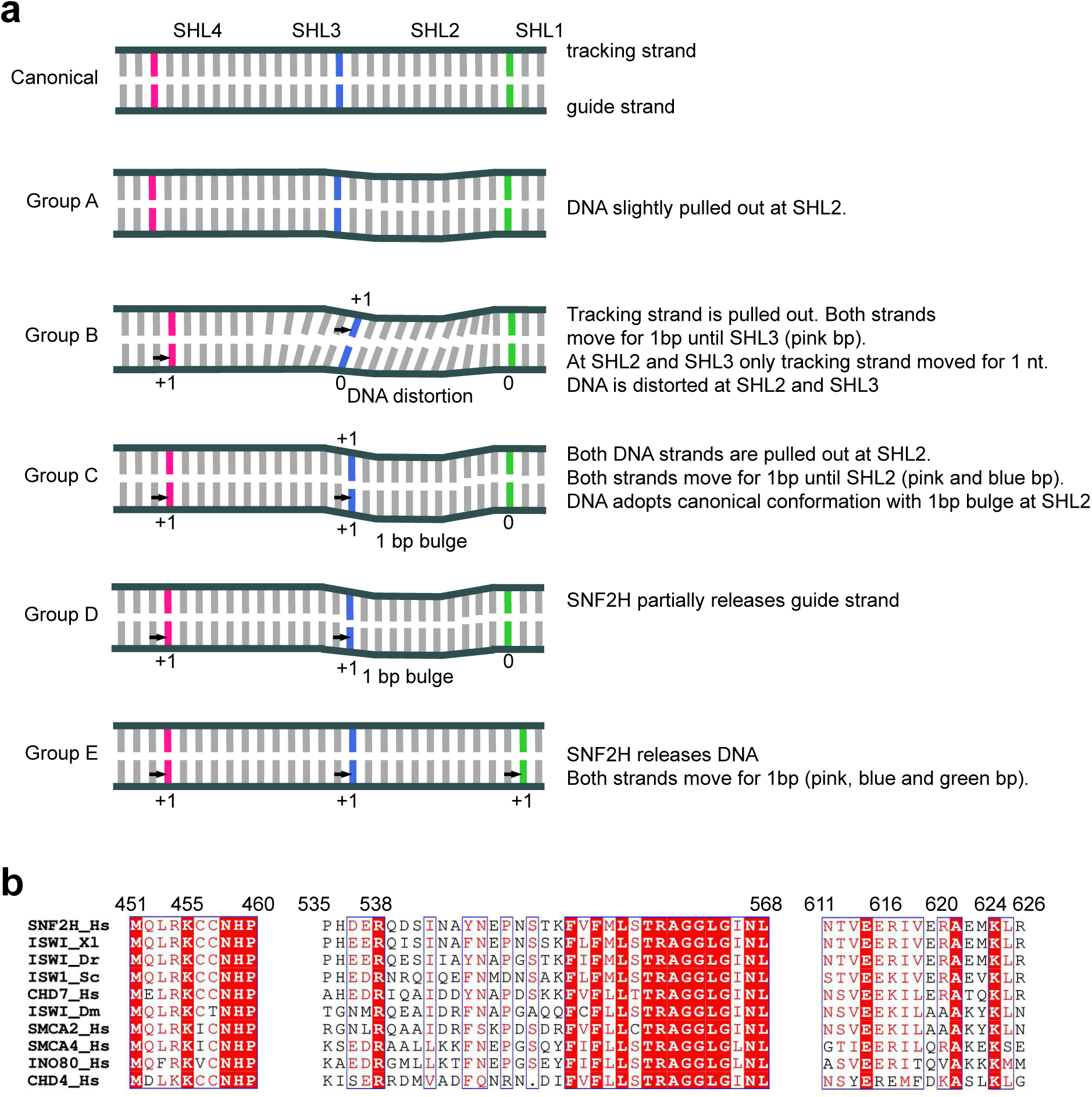
Model for DNA translocation on the nucleosome. **a)** Schematic model showing DNA translocation on the nucleosome by SNF2H. **b)** Sequence alignment of motor enzymes from different chromatin remodeling families, showing conservation of key residues that interact with DNA.

**Supplementary Table S1:**
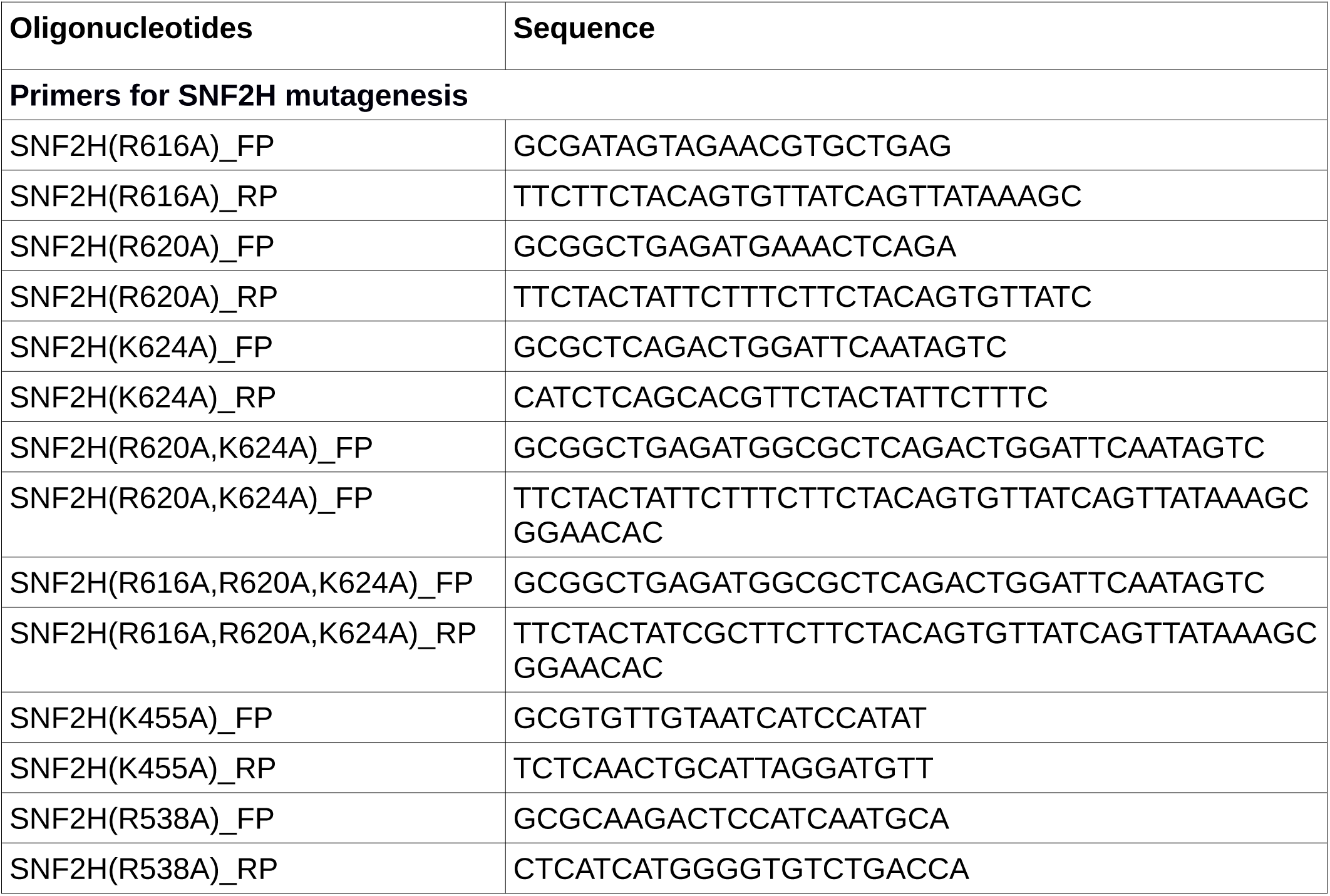
List of oligonucleotides used in the study.

**Supplementary Table S2:** Plasmids used in this study.

|  |  |
| --- | --- |
| SJ559 | SNF2H(R616A) |
| SJ560 | SNF2H(R620A) |
| SJ561 | SNF2H(K624A) |
| SJ562 | SNF2H(R620A,K624A) |
| SJ563 | SNF2H(R616A,R620A,K624A) |
| SJ564 | SNF2H(K455A) |
| SJ565 | SNF2H(R538A) |
| SJ566 | SNF2H(K455A,R538A) |
| SJ186 | SNF2H(WT) |
| SJ298 | 601 widom sequence in pET3a vector |

## Notes

### Competing Interest Statement

The authors have declared no competing interest.

